# Metabolomic signature of angiopoietin-like protein 3 deficiency in fasting and postprandial state

**DOI:** 10.1101/441857

**Authors:** Emmi Tikkanen, Ilenia Minicocci, Jenni Hällfors, Alessia Di Costanzo, Laura D’Erasmo, Eleonora Poggiogalle, Lorenzo Maria Donini, Peter Würtz, Matti Jauhiainen, Vesa M. Olkkonen, Marcello Arca

**Affiliations:** Nightingale Health Ltd., Helsinki, Finland; Department of Internal Medicine and Medical Specialties, Sapienza University of Rome, Italy; Department of Experimental Medicine, Sapienza University of Rome, Italy; Minerva Foundation Institute for Medical Research, Biomedicum 2U, Helsinki, Finland; Department of Anatomy, Faculty of Medicine, University of Helsinki, Finland

**Keywords:** Metabolomics, biomarkers, ANGPTL3, drug target, lipoproteins, lipid lowering

## Abstract

**Objective:** Loss-of-function variants in the angiopoietin-like 3 gene (*ANGPTL3*) have been associated with low levels of plasma lipoproteins and decreased coronary artery disease risk. We aimed to determine detailed metabolic effects of genetically-induced ANGPTL3 deficiency in fasting and postprandial state.

**Approach and Results:** We studied individuals carrying S17X loss-of-function mutation in *ANGPTL3* (6 homozygous and 32 heterozygous carriers) and 38 noncarriers. Nuclear magnetic resonance metabolomics was used to quantify 225 circulating metabolic measures. We compared metabolic differences between loss-of-function carriers and noncarriers in fasting state and after a high fat meal. In fasting, ANGPTL3 deficiency was characterized by similar extent of reductions in low-density lipoprotein cholesterol (0.74 SD-units lower concentration per loss-of-function allele [95%CI 0.42–1.06]) as observed for many triglyceride-rich lipoprotein measures, including very-low-density lipoprotein cholesterol (0.75 [0.45–1.05]). Within most lipoprotein subclasses, absolute levels of cholesterol were decreased more than triglycerides, resulting in the relative proportion of cholesterol being reduced within triglyceride-rich lipoproteins and their remnants. Further, beta-hydroxybutyrate was elevated (0.55 [0.21–0.89]). Homozygous *ANGPTL3* loss-of-function carriers showed essentially no postprandial increase in triglyceride-rich lipoproteins and fatty acids, without evidence for adverse compensatory metabolic effects.

**Conclusions:** In addition to overall triglyceride and low-density lipoprotein cholesterol lowering effects, ANGPTL3 deficiency results in reduction of cholesterol proportion within triglyceride-rich lipoproteins and their remnants. Further, *ANGPTL3* loss-of-function carriers had elevated ketone body production, suggesting enhanced hepatic fatty acid beta-oxidation. The detailed metabolic profile in human knockouts of *ANGPTL3* reinforces inactivation of ANGPTL3 as a promising therapeutic target for decreasing cardiovascular risk.

**HIGHLIGHTS:**

- ANGPTL3 deficiency results in similar reductions in LDL cholesterol and many triglyceride-rich lipoprotein lipids measures, such as VLDL cholesterol, with no evidence of substantial adverse effects on the comprehensive panel of circulating metabolite biomarkers tested here.
- In particular, ANGPTL3 deficiency results in reduction of cholesterol content in triglyceride-rich lipoproteins and their remnants, which have been highlighted as risk factor for cardiovascular disease independently of LDL levels.
- Homozygous *ANGPTL3* loss-of-function carriers show essentially no postprandial increase in triglyceride-rich lipoproteins and fatty acids in response to a fat challenge, and display consistently elevated postprandial levels of ketone bodies and lactate when compared to noncarriers, suggesting enhanced hepatic fatty acid beta-oxidation.

## INTRODUCTION

Considerable risk for cardiovascular disease (CVD) persists in individuals treated with low-density lipoprotein (LDL) cholesterol lowering statin and PCSK9 inhibitor therapies^1^. Experimental models and human genetics have indicated that triglyceride-rich lipoproteins (TRL) play a causal role in the pathogenesis of CVD^2^. Post-hoc analyses of statin trials have further shown that reductions of TRL is associated with lower cardiovascular event risk independently of the LDL cholesterol reduction achieved from statins^3,4^. These observations have accelerated the development of novel TRL-lowering therapeutics for CVD prevention, with drug targets informed by recent discoveries from genetic studies^5^. These include identification of loss-of-function (LOF) mutations in the angiopoietin-like 3 gene (*ANGPTL3*) causing familial combined hypolipidemia (FHBL2; OMIM #605019)^6,7^. This Mendelian condition is characterized by simultaneous presentation of low circulating concentrations of triglycerides, LDL cholesterol and high-density lipoprotein (HDL) cholesterol levels, as well as potential beneficial effects on glucose metabolism^7^.

ANGPTL3 is a protein secreted by the liver, and its deficiency was first identified in a hypolipidemic mouse strain^8^. The ANGPTL3 protein is as a potent inhibitor of lipoprotein lipase, a primary factor that clears TRL from the circulation^9^. Reduced ANGPTL3 also reduce hepatic apolipoprotein (apo) B secretion and increased hepatic LDL uptake, leading to reduced plasma LDL cholesterol levels^10^. ANGPTL3 further acts to inhibit endothelial lipase, which may contribute to the low HDL cholesterol levels in *ANGPTL3* LOF carriers^11^. The first human *ANGPTL3* LOF mutations described were nonsense mutations S17X and E129X in the first exon of the gene^6^. Since the initial publications on these mutations, several other *ANGPTL3* LOF mutations have been characterized^12-14^. Individuals with complete ANGPTL3 deficiency have been reported to lack significant coronary atherosclerotic plaques^15^. Exome sequencing of large epidemiological cohorts have further shown that heterozygous carriers of *ANGPTL3* LOF variants have a 35–40% reduced risk of coronary artery disease compared to the general population^15,16^. Moreover, no increased prevalence in fatty liver disease, or other apparent adverse health effects have been identified for *ANGPTL3* LOF carriers, including homozygote carriers with complete ANGPTL3 deficiency^7,17^. Thus, ANGPTL3 is a promising therapeutic target for lowering CVD risk, and clinical trials of monoclonal antibodies or antisense oligonucleotides for ANGPTL3 inhibition have recently shown to lead to substantial lowering of circulating triglycerides and LDL cholesterol levels.^16,18^

Despite the promising results from phase 1 trials targeting ANGPTL3^16,18^, a number of important questions of the molecular effects of ANGPTL3 inhibition remain. First, it is unclear which specific lipid components of TRL are mostly affected by ANGPTL3 deficiency, and thus the underlying mechanism of reduced CVD risk is incomplete understood. Second, the effects of ANGPTL3 deficiency on many emerging biomarkers for cardiometabolic risk have not been addressed^9^. We have previously shown that nuclear magnetic resonance (NMR) metabolomics is a powerful method to characterize the fine-grained effects of *PCSK9* and *HMGCR* genetic variants on a lipid metabolism and other metabolic pathways, with a close match to the changes observed in statin trials^19,20^. Here, we use NMR metabolomics to characterize the systemic effects of ANGPTL3 deficiency on detailed measures of lipoprotein composition, fatty acids, and circulating metabolites. Because most people are in a non-fasting state for the majority of the day, studying the effects on postprandial metabolism may further contribute to explain the cardioprotective mechanism of ANGPTL3 deficiency. We have previously reported highly reduced postprandial response in triglycerides among *ANGPTL3* LOF homozygotes^21^. Here we further examined the detailed metabolic effects of ANGPTL3 deficiency after a high fat meal.

## MATERIAL AND METHODS

### Study cohort of *ANGPTL3* loss-of-function carriers

An overview of the study design in shown in the “Graphic summary”. The study population and the design of the oral fat tolerance challenge have been described in detail elsewhere^21^. Briefly, 6 homozygous and 32 heterozygous carriers of *ANGPTL3* S17X LOF mutation (henceforth *ANGPTL3* LOF) and 38 non-carriers were considered in the present study. Clinical examinations and blood sample drawings were performed after an overnight fast, after which participants underwent an oral fat tolerance test. The test meal (muffin prepared with olive oil, eggs, ricotta cheese, nuts, cocoa, wheat flour, and skimmed milk) consisted of 73 g fat, 52 g carbohydrate, 22 g protein, and 145 mg cholesterol. Blood samples were drawn before the test meal and at 2, 4, and 6 hours after the meal. The Ethical Committee of Sapienza University of Rome approved the study protocol and all study participants provided their informed consent.

### Lipid and metabolite quantification

Frozen EDTA plasma samples from fasting state and those collected during the oral fat tolerance test were used for metabolomic analyses. Metabolic biomarkers were quantified using proton NMR metabolomics (Nightingale Health Ltd, Helsinki, Finland). This method provides simultaneous quantification of routine lipids, lipoprotein subclass profiling with lipid concentrations within 14 subclasses, fatty acid composition, and various low-molecular metabolites including amino acids, ketone bodies and gluconeogenesis-related metabolites in molar concentration units. Details of the experimentation and applications of the NMR metabolomics platform have been described previously^19,20,22,23^.

### Statistical analysis

We first evaluated associations between *ANGPTL3* LOF carrier status and fasting metabolite measures. Prior to analyses, the concentrations of all metabolic markers were log-transformed and scaled to standard deviation (SD) units to enable comparison of results for measures with different units and across wide ranges of concentrations. As the primary analysis, we examined an additive model for association between metabolites and the three genotype classes (*ANGPTL3* LOF homozygotes, heterozygotes, and noncarriers). For each metabolic measure, the concentration difference (in log-transformed and subsequently SD-scaled units) per *ANGPTL3* LOF allele was calculated using linear regression adjusted for age and gender. As a secondary analysis, we studied the associations of complete *ANGPTL3* knockouts and fasting metabolites using a recessive model (homozygous carriers vs. heterozygotes and noncarriers combined). Due to the correlated nature of metabolomics measures, significance level was defined to be 0.05/23, where 23 is the number of principal components explaining 99% of the variation of all metabolic measures at fasting state.

To evaluate postprandial responses in metabolites, we calculated means and standard errors by *ANGPTL3* LOF carrier status at each time point. Postprandial dynamics in the three genotype classes were compared using net incremental area under the curve (iAUC) statistics (AUC that differs from the fasting value) with the trapezoidal rule^24^. The differences in iAUCs between the three genotype classes were tested with *t*-tests.

## RESULTS

### Study participants

The clinical characteristics of the study participants are summarized in **Table 1**. *ANGPTL3* LOF carriers and noncarriers were comparable for age, gender, body mass index, and waist-hip-ratio. Plasma levels of ANGPTL3 protein were undetectable in homozygotes, while heterozygotes had 51% lower concentration compared to that of noncarriers. The three genotype classes were comparable for dietary intake, physical activity, smoking prevalence and use of anti-inflammatory medications^21^. Descriptive statistics for all metabolic biomarkers tested are reported in **Supplementary Table 1**.

**Table 1.** Characteristics of study participants.

|  | **Noncarriers** | ***ANGPTL3***<br>**Heterozygous**<br>**Carriers** | ***ANGPTL3***<br>**Homozygous**<br>**Carriers** |
| --- | --- | --- | --- |
| **N** | 38 | 32 | 6 |
| **Age, years** | 47.0 (12.6) | 50.8 (14.5) | 53.5 (24.5) |
| **Gender, females** | 21 (55.3) | 15 (47.0) | 4 (66.7) |
| **Body mass index (kg/m<sup>2</sup>)** | 26.3 (4.2) | 28.8 (4.4) | 27.8 (5.0) |
| **Statin use** | 1 (2.6) | 2 (6.3) | 0 |
| **Fasting *ANGPTL3* protein**<br>**concentration (ng/ml)** | 208.7 (80.3) | 108.3 (60.1) | 0 |
| Data are presented as mean (SD) or number of individuals (%) |  |  |  |

### Effects of *ANGPTL3* loss-of-function on lipids and metabolites at fasting

Compared with noncarriers, carriers of *ANGPTL3* LOF mutations had substantially reduced levels in almost all lipid measures assayed by NMR metabolomics. These include lower concentrations of routine lipid measures, as well as the cholesterol and triglyceride levels in major subfractions and reduced concentrations of apolipoproteins B and A1 (**Figure 1, Supplementary Table 2)**. Scaled to the same variation in each lipid measure (i.e. in units of SD), we observed comparable extent of lowering effects on LDL cholesterol levels (−0.74 SD-units of concentration per allele, [95% confidence interval −1.06 to −0.42]) as observed for VLDL cholesterol levels (−0.75, 95% CI −1.05 to −0.45) and other measures of TRL. The lowering effect size on total plasma triglycerides was −0.62 (95% CI −0.93 to −0.32). In absolute concentrations, these lowering effects correspond to 0.28 mmol/L lower for LDL cholesterol, 0.13 mmol/L lower for VLDL cholesterol, and 0.20 mmol/L for total triglycerides. Also HDL cholesterol and other lipid measures in HDL particles were lowered to a similar extend when scaled to SD-concentrations of same variation. The effects of *ANGPTL3* LOF on all tested metabolites are shown in **Supplementary Figures 1-2**.

**Figure 1.**
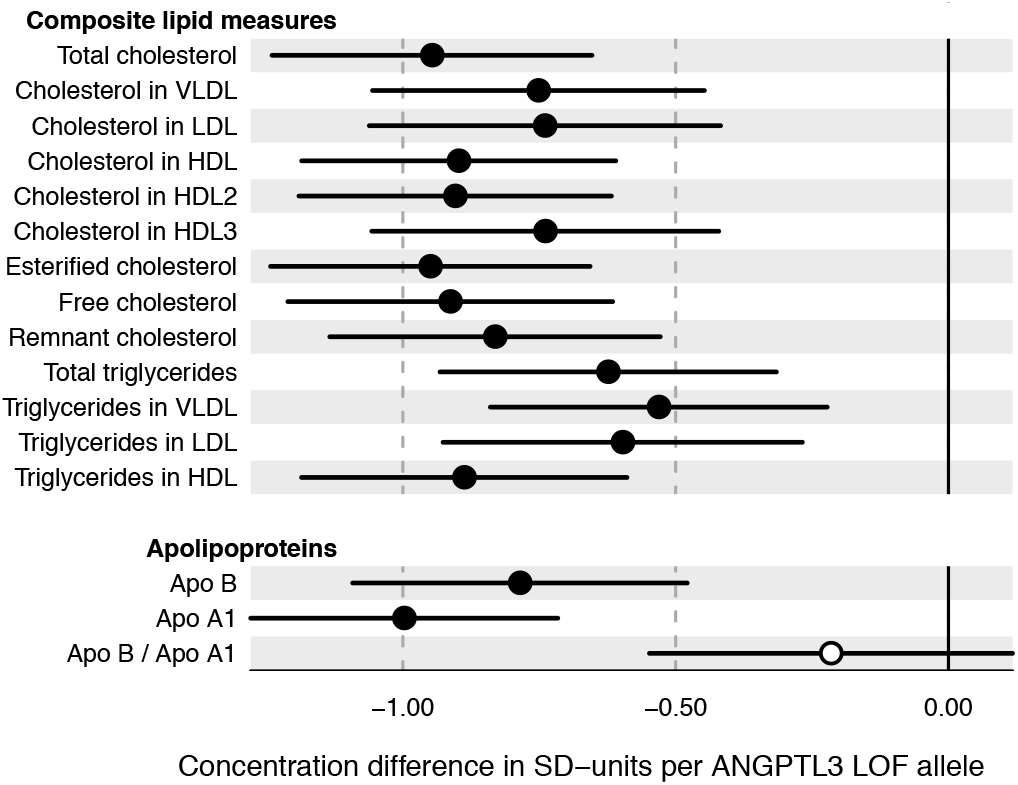
Effects of *ANGPTL3* loss-of-function variant on fasting lipoprotein lipids and apolipoproteins. Effect estimates are shown as difference in SD-scaled concentration units per *ANGPTL3* S17X allele (additive model). Error bars indicate 95% confidence intervals. Filled and open circles denote P-value for association below and above 0.002, respectively.

Zooming in on the more specific effects of *ANGPTL3* LOF on lipoprotein subclass measures, we observed substantial reduction of cholesterol concentration within all lipoprotein subclasses (**Figure 2A**). The lowering effects were somewhat stronger for intermediate-density lipoprotein (IDL) in comparison to other apoB-100 containing particles. For HDL subclasses, cholesterol levels were most prominently lowered in medium-sized HDL particles. Scaled to the same variation, the extent of cholesterol lowering in all 10 apoB-containing subclasses was broadly similar to that observed for the routine lipid measures. Analogously, the triglyceride concentration in all lipoprotein subclasses (except for small HDL) were reduced; however the triglyceride lowering was to a somewhat smaller extent for IDL and other similarly-sized lipoprotein particles than that observed for cholesterol. Thus, even though the absolute triglyceride levels in lipoprotein particles were reduced, the relative proportion of triglycerides in TRL particles was actually increased due to ANGPTL3 deficiency. Concordantly, the cholesterol proportion of medium-sized and small VLDL and IDL particles decreased (**Figure 2A** lower panels). This is further illustrated in **Figure 2B**, which depicts the overall lipid composition for selected lipoprotein subclasses in *ANGPTL3* LOF carriers and noncarriers. For example, the proportion of cholesterol in small VLDL was 36% in noncarriers, 34% in *ANGPTL3* LOF heterozygotes and 26% in homozygotes. The cholesterol proportion was also reduced in large LDL (67%, 66%, and 62%, respectively), whereas in large HDL, *ANGPTL3* LOF homozygotes had the highest proportion of cholesterol (55% vs. 47% and 46% in *ANGPTL3* LOF heterozygotes and noncarriers, respectively).

**Figure 2.**
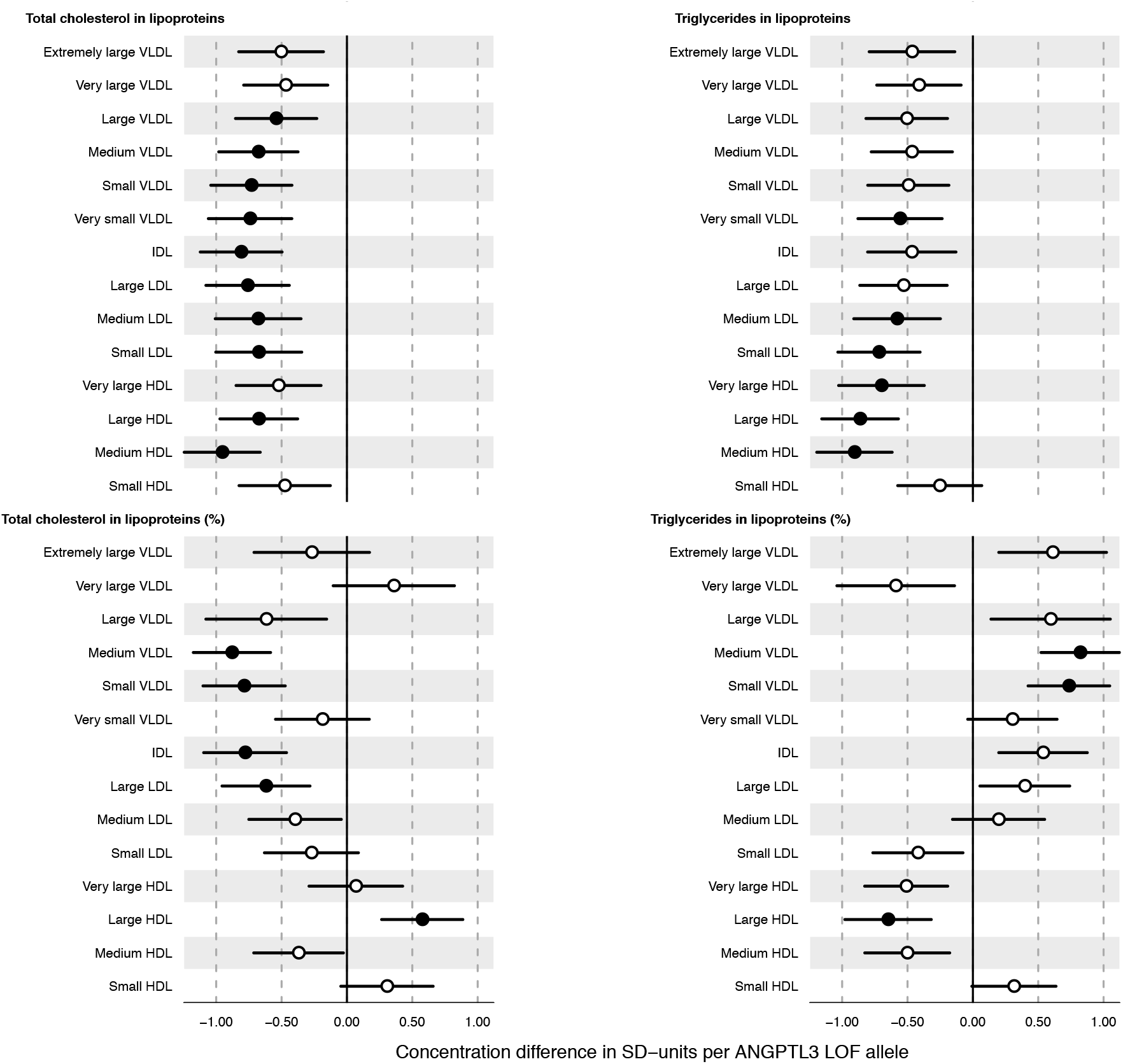
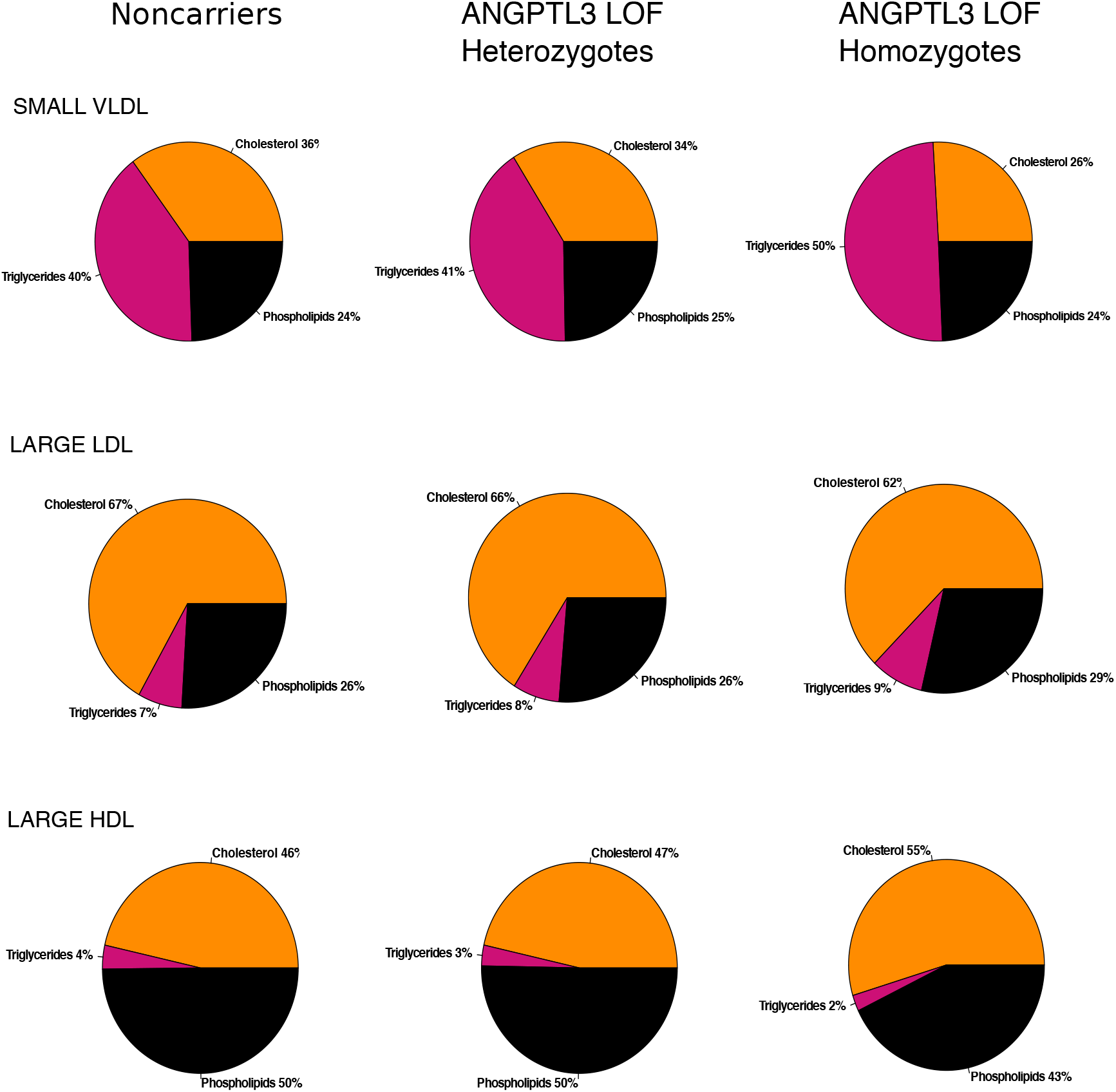
Effects of *ANGPTL3* loss-of-function variant on fasting cholesterol and triglycerides concentrations in lipoprotein subclasses and their composition. **A)** Effect estimates are shown as difference in SD-scaled concentration units per *ANGPTL3* S17X allele (additive model). Error bars indicate 95% confidence intervals. Filled and open circles denote P-value for association below and above 0.002, respectively. **B)** Average lipid content of selected lipoprotein subclasses in *ANGPTL3* LOF carriers and noncarriers are depicted to further illustrate the effects on lipoprotein composition.

The substantial lowering effects of ANGPTL3 deficiency on lipoprotein lipids may influence both absolute concentrations of circulating fatty acid and their relative proportions. We found that *ANGPTL3* LOF carriers displayed a significant reduction of total fatty acids, including saturated, monounsaturated, and polyunsaturated omega-3 and omega-6 fatty acids (**Figure 3**). Assessment of the proportions of fatty acid classes (i.e. their ratios relative to total fatty acids) indicated an elevation of the proportion of saturated fatty acids and a reduction in the proportion of omega-3 fatty acids.

To assess potential nonlipid effects of ANGPTL3 deficiency, we examined the changes in circulating amino acids, glycolysis and gluconeogenesis substrates and products, ketone bodies, and other metabolites quantified simultaneously alongside lipid measures by the NMR metabolomics platform. Several of these nonlipid biomarkers also showed significant association with *ANGPTL3* LOF carrier status (**Figure 3**). The ketone body beta-hydroxybutyrate, a biomarker of hepatic fatty acid beta-oxidation, was markedly elevated in *ANGPTL3* LOF carriers (0.55 SD per allele, 95% CI 0.21 to 0.89). Further, the energy metabolism intermediate acetate was reduced (−0.56 SD per allele, 95% CI −0.90 to −0.22). Citrate levels also displayed a similar tendency. Branched-chained and aromatic amino acids, which are biomarkers for increased type 2 diabetes and CVD risk^23,25^, did not differ between *ANGPTL* LOF carriers and noncarriers. Further, we observed no robust differences in creatinine, an indicator of kidney function, or in glycoprotein acetyls (GlycA) level, a marker of chronic inflammation.

**Figure 3.**
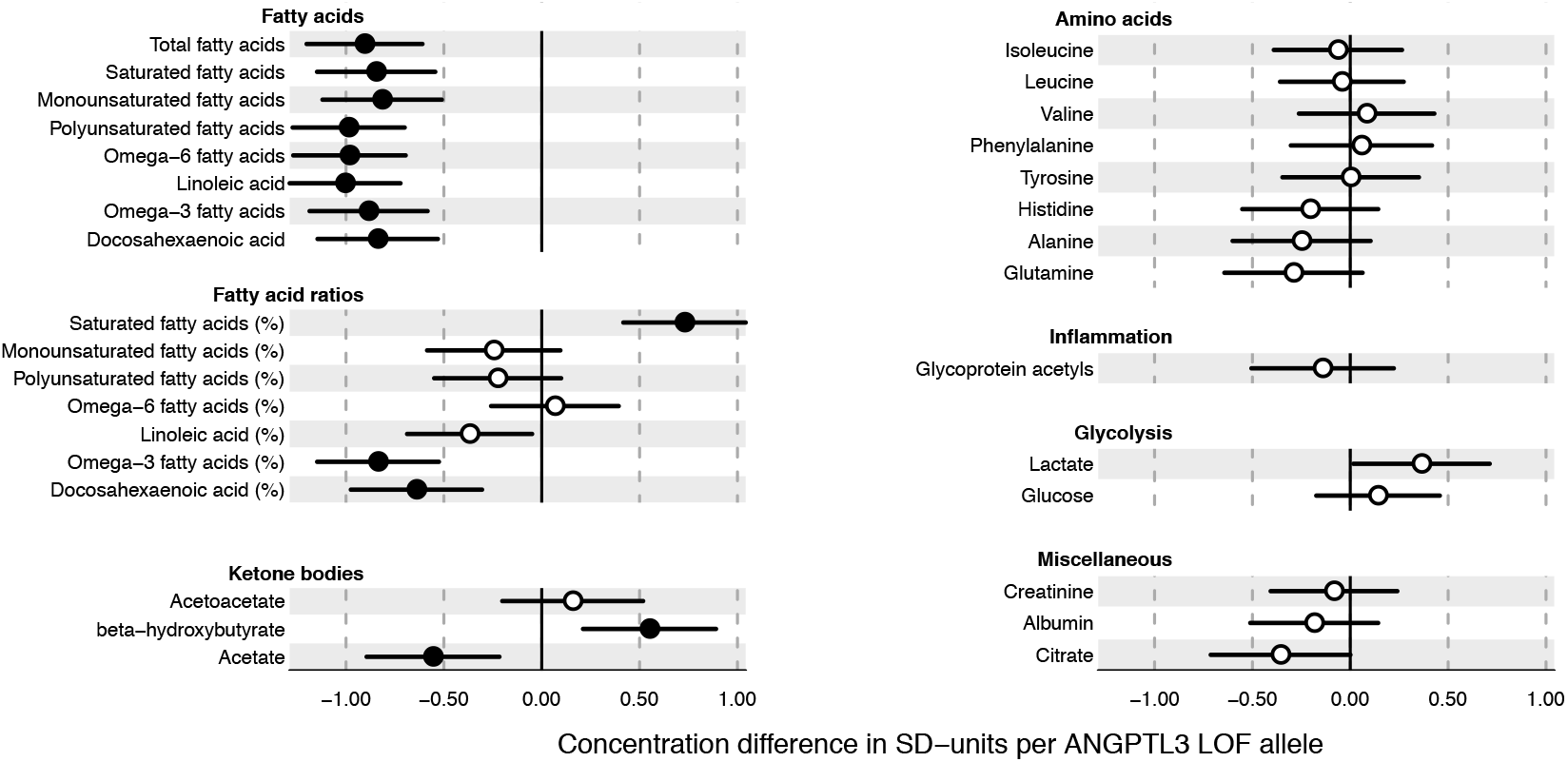
Effects of *ANGPTL3* loss-of-function variant on fatty acids and polar metabolites. Effect estimates are shown as difference in SD-scaled concentration units per *ANGPTL3* S17X allele (additive model). Error bars indicate 95% confidence intervals. Filled and open circles denote P-value for association below and above 0.002, respectively.

In analyses employing a recessive model there was evidence for individuals with complete ANGPTL3 deficiency having more than twice the lipid lowering effects of heterozygote *ANGPTL3* LOF carriers. For example, homozygous *ANGPTL3* LOF carriers had on the average −1.97 (95% CI −2.69 to −1.25) SD units lower LDL cholesterol, and −1.86 (95% CI −2.56 to −1.16) lower VLDL cholesterol, when compared to heterozygous *ANGPTL*3 LOF carriers and noncarriers. Many other lipids and lipoprotein measures also had substantially larger effect sizes for homozygous *ANGPTL3* LOF carriers in a recessive model (**Supplementary Figures 3–4, Supplementary Table 3**). None of the non-lipid markers were significant in the recessive model or displayed evidence for deviation from an additive model (**Supplementary Figures 3– 4, Supplementary Table 3**).

### Effects of ANGPTL3 loss-of-function on metabolic response to a fat challenge

To examine the effects of ANGPTL3 deficiency on postprandial metabolism, we further assessed the lipoprotein and metabolite trajectories after an oral fat challenge. Postprandial responses in selected metabolic measures for *ANGPTL3* LOF carriers and noncarriers are shown in **Figure 4**. *ANGPTL3* LOF homozygotes had drastically lower concentration of cholesterol in VLDL, LDL and IDL particles in all timepoints in comparison to *ANGPTL3* LOF heterozygotes and noncarriers. There was essentially no change in VLDL cholesterol after the fat challenge among *ANGPTL3* LOF homozygotes, whereas *ANGPTL3* LOF heterozygotes and noncarriers showed substantially increased levels (P_iAUC_ = 0.07 for difference in iAUC). This effect on VLDL cholesterol seemed to be driven particularly by differences for large and medium sized VLDL particles in response to the high fat meal. For instance, both *ANGPTL3* LOF heterozygotes and non-carriers displayed more than 80% increase in cholesterol in large VLDL at four hours after the meal when compared to *ANGPTL3* LOF homozygotes (P_iAUC_=0.02 for *ANGPTL3* LOF homozygotes versus noncarriers). In contrast, the high fat meal did not give rise to an increase in cholesterol levels of small VLDL, IDL and LDL particles for any of the groups; rather, there was a small decrease in cholesterol within these smaller apoB-carrying lipoprotein particles.

**Figure 4.**
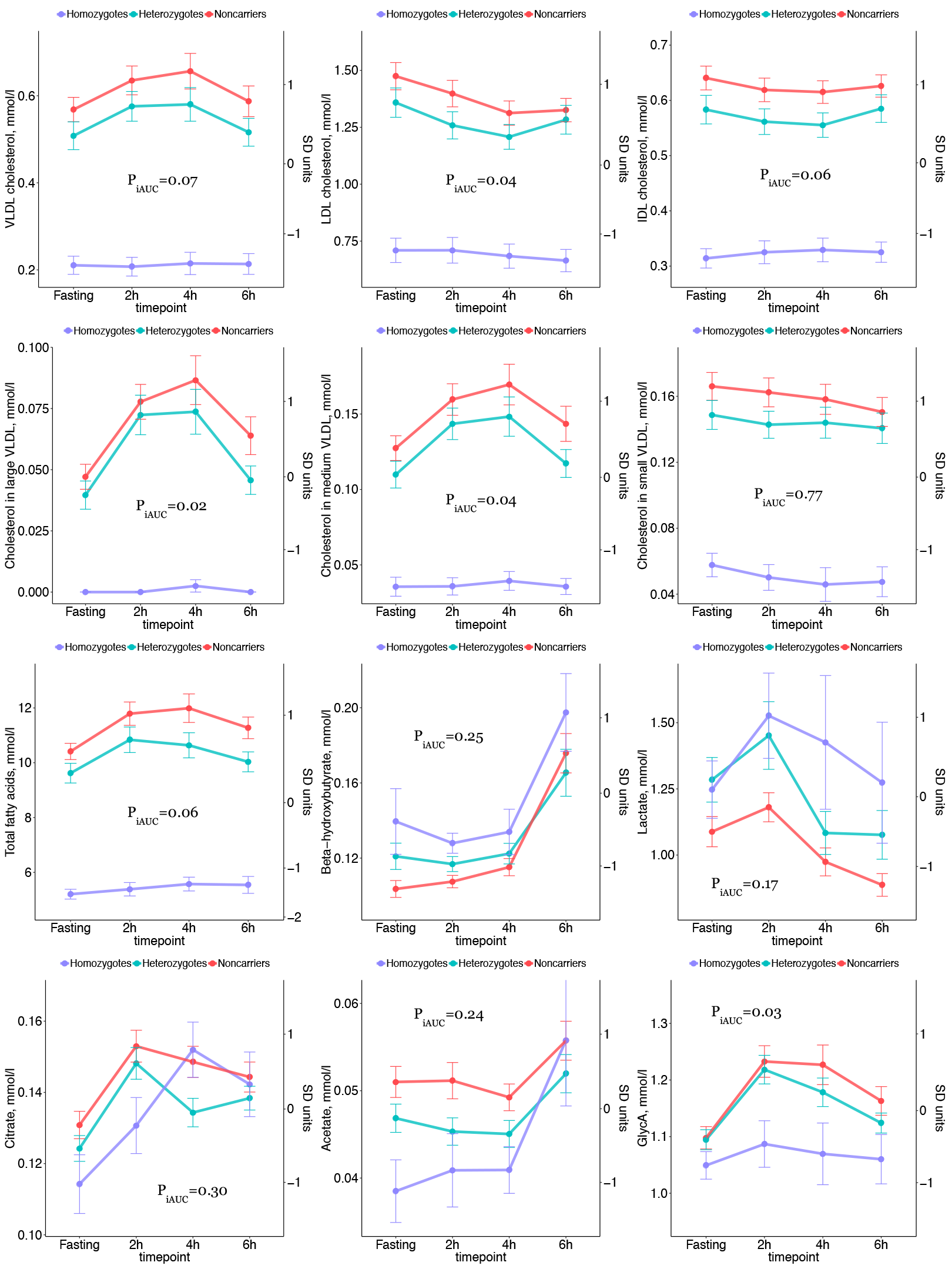
Effects of *ANGPTL3* loss-of-function carrier status on metabolic response to an oral fat tolerance test. Results are shown as mean concentration and standard errors at fasting and postprandial blood sample draw for selected biomarkers. Left hand side axes indicate concentrations in absolute units and right side indicates concentations in SD-scaled units. P_iAUC_ indicates significance of difference in incremental area under the curve, reflecting meal response for ANGPTL3 S17X loss-of-function homozygotes versus noncarriers.

The postprandial response to the oral fat challenge in total fatty acids, beta-hydroxybutyrate and lactate are shown in **Figure 4**. Homozygous *ANGPTL3* LOF carriers had markedly lower total fatty acids at all time points in comparison to the other two groups, and showed essentially no postprandial increase in fatty acid concentrations. While the absolute concentration of omega-3 fatty acids was essentially unaffected by the fat challenge, the proportion of omega-3 fatty acids to total fatty acids increased after the meal in *ANGPTL3* LOF homozygotes compared to other groups (P_iAUC_=0.0003, **Supplementary Figure 5**). The responses in absolute levels of saturated and monounsaturated fatty acids followed similar patterns as that of total fatty acids, with essentially no postprandial response for *ANGPTL3* LOF homozygotes; however, the relative proportion of monounsaturated fatty acids increased for all groups (**Supplementary Figure 5**). The shape of the postprandial response in ketone bodies was broadly similar for *ANGPTL3* LOF carriers and noncarriers, but the concentrations of beta-hydroxybutyrate were maintained elevated in the homozygous *ANGPTL3* LOF carriers (**Figure 4**). *ANGPTL3* LOF carriers also displayed consistently higher lactate levels after the fat meal challenge, and increases in postprandial citrate and acetate levels. Among other non-lipid biomarkers, glycoprotein acetyl (GlycA; a marker of chronic inflammation) levels showed a blunted postprandial response in *ANGPTL3* LOF carriers, possibly indicating a smaller inflammatory effect of the high fat meal (P_iAUC_=0.03, **Figure 4**). Postprandial responses for other selected biomarkers are shown in **Supplementary Figure 6**.

## DISCUSSION

In the present work we describe a detailed metabolic signature associated with ANGPTL3 deficiency in humans. Because ANGPTL3 deficiency is known to influence postprandial lipid metabolism via modulation of lipoprotein lipase activity^21^, we considered it particularly interesting to investigate the fine-grained metabolic effects of *ANGPTL3* LOF also after a high-fat meal. We found that ANGPTL3 deficiency is characterized by markedly low levels of fasting and postprandial LDL and VLDL cholesterol, as well as other measures of TRL. In particular, individuals with two inactivating mutations in *ANGPTL3* and no measurable concentration of ANGPTL3 protein showed essentially no increase in TRL in response to a high fat meal. The very low lipid levels among these homozygote carriers, who effectively act as human knockouts for ANGPTL3, were maintained after the oral fat challenge without evidence for adverse compensatory effects on the extensive metabolic biomarker panel examined in this study.

The main findings of our study are 4-fold: First, we found that ANGPTL3 deficiency results in similar reductions in magnitude for LDL and VLDL cholesterol, when scaled to the same variation in each lipid measure. Thus, ANGPTL3 inhibitors should be more effective in lowering VLDL cholesterol as compared to statins or PCSK9 inhibitors^19,20^, suggesting added benefit of ANGPTL3 inactivating therapy in individuals with combined dyslipidemias. This finding has clinical relevance, because increasing evidence suggest that the cholesterol content of VLDL and other TRL are causally implicated in the development of CVD, independent of LDL cholesterol^2,3,26^. Although TRL have been suggested to play a causal role in CVD in Mendelian randomization studies^27^, the triglycerides *per se* are unlikely to be the causal factor because these lipids are not accumulating in the atherosclerotic plaque. In contrast, the cholesterol within TRL and their remnants can accumulate in the arterial intima, and even get trapped more easily than LDL cholesterol^2^. In line with our results, a recent phase 1 trial on antisense oligonucleotides targeting hepatic *ANGPTL3* messenger RNA has reported up to 60% reduction in VLDL cholesterol and 33% reduction in LDL cholesterol^18^.

Second, the detailed profiling of lipoprotein subclasses revealed that the absolute concentrations of both cholesterol and triglycerides in most lipoprotein subclasses were reduced in *ANGPTL3* LOF carries compared to noncarriers. However, the proportion of cholesterol (relative to the total lipid concentration in a given subclass) was reduced and the proportion of triglycerides increased in small and medium-sized VLDL and IDL subclasses. These results indicate that ANGPTL3 deficiency leads to less cholesterol load in the composition of TRL, which could potentially contribute to explain the lower CVD risk observed among *ANGPTL3* LOF carriers. We observed no clear differences in the patterns of postprandial response of TRL measures for *ANGPTL3* LOF heterozygotes as compared to noncarriers. In contrast, there was a remarkable lack of increase in TRL after the fatty meal intake among the 6 individuals with complete ANGPTL3 deficiency, as observed previously with more narrow measures of lipid metabolism^21,28^. This lack of a dose-dependent postprandial response could suggest that therapeutic inhibition of ANGPTL3 may not alleviate the increase in TRL after a meal intake unless protein levels of ANGPTL3 are lowered to a very high extent. To this regard, we have recently determined that plasma ANGPTL3 less than 60 ng/ml represents a critical threshold associated with marked reductions in fasting lipid and lipoprotein concentrations^29^. Initial trial results have demonstrated therapeutic reductions of plasma ANGPTL3 concentration of up to 85%^18^. Further studies are required to elucidate whether this extent of ANGPTL3 protein depletion is sufficient to substantially blunt the postprandial response in TRL or whether complete ANGPTL3 deficiency is required to observe this.

Third, we found that ANGPTL3 deficiency causes an elevation of fasting concentrations of the ketone beta-hydroxybutyrate, which was maintained throughout the postprandial response. During ketogenesis in the liver, fatty acids are converted into ketone bodies via beta-oxidation to produce energy. As ketogenesis is elevated when the influx of fatty acids into the liver is increased^30^, the present observation may indicate that complete ANGPTL3 deficiency is accompanied by an increased hepatic utilization of fatty acids derived from the lipolysis of TRL or from adipose tissue. However, since the shape of postprandial response in beta-hydroxybutyrate and the other ketone body measured, acetoacetate, was similar independent of the ANGPTL3 carrier status, one could speculate that this phenomenon might be related to partitioning of adipose tissue-derived fatty acid during the interprandial phase. However, this issue requires further investigations. Similarly to beta-hydroxybutyrate, lactate levels were consistently higher at fasting and after the meal challenge for complete ANGPTL3 deficiency. This may indicate an enhanced conversion of pyruvate to lactate instead of its routing to the acetate pathway, consistent with the reduced concentration of acetate in the LOF carriers. These observations may suggest that ANGPTL3 deficiency is causing a modest shift of energy substrate utilization. The potential health effects of enhanced ketone body production remain unclear, but large epidemiological studies have generally reported only weak or null associations of ketone bodies with cardiometabolic risk factors and disease outcomes^22^.

A further insight from this study arises from the metabolic biomarkers not associated with ANGPTL3 deficiency, in particular for the homozygous carriers. These null results suggest there are no substantial adverse effects of complete absence of ANGPTL3 protein on amino acids and glycolysis-related metabolites, neither at fasting nor as a compensatory mechanism for the absence of postprandial lipid increase. This is informative for the safety of ANGPTL3 inhibiting therapeutics, because many of these metabolites have been shown to be biomarkers of the risk for type 2 diabetes and CVD events^23,25^. This is also the case for several of the relative proportions of fatty acid analyzed in this study. The observed modulations in the fatty acid balance for the relative proportions of saturated and omega-3 fatty acids could potentially be unfavorable in terms of cardiovascular risk, but the causal relations of these measures remain unclear. For the postprandial effects, it is noteworthy that the inflammatory biomarker glycoprotein acetyls was low in *ANGPTL3* LOF homozygotes, and less elevated in the postprandial stage; such blunted increase in low-grade inflammation in response to a fatty meal might contribute to additional risk-reducing effects beyond lipids. Overall, these results illustrate the potential of wider metabolic profiling of human knockouts for drug targets for elucidating molecular mechanisms and potential metabolic side effects.

The strengths of the present study include a unique setting with fasting and postprandial measurements in *ANGPTL3* LOF carriers, including the rare instance of six individuals with complete ANGPTL3 deficiency. We used NMR metabolomics to obtain detailed measures of lipid metabolism and non-lipid biomarkers from multiple metabolic pathways. However, we acknowledge that other metabolomics assays capable of capturing additional blood metabolites could further contribute to characterize the metabolic effects beyond lipid metabolism. Despite the large effects on many lipid levels caused by ANGPTL3 deficiency, the small sample size provided limited power to detect all relevant metabolic associations, in particular for robustly comparing differences in the postprandial response. Our study was limited to characterize carriers of *ANGPTL3* S17X LOF mutation only. Thus, there is a need for further metabolic characterization of other LOF variants in the *ANGPTL3* gene. Detailed metabolic profiling in large biobanks with sequencing information available provides an opportunity to enable this and further extend applications to other lipid-lowering targets^20,31^.

In conclusion, this detailed metabolic profiling study demonstrates that ANGPTL3 deficiency is characterized by substantial reductions in VLDL particles and their remnants, and a lower cholesterol content in these lipoproteins. Further, complete ANGPTL3 deficiency leads to virtual absence of postprandial increase in TRL. These findings support the increasing body of evidence indicating that genetic inhibition of ANGPTL3 causes a broad range of beneficial lipid changes, without adverse compensatory metabolic effects. Detailed metabolic profiling in trials of pharmacologic inactivation of ANGPTL3 could further help to confirm these findings in clinical settings and, in combination with the genetic evidence, uncover potential off-target effects^32^.

## ACKNOWLEDGEMENTS

Study participants are acknowledged for their availability and commitment. The authors thank Jari Metso, M.Sc. for excellent laboratory assistance and Anna Montali, B.Sc., for her assistance in recruiting and following-up study participants.

